# LRSPAT: A low-rank framework for spatial omics statistics

**DOI:** 10.64898/2026.08.26.747291

**Authors:** H. Robert Frost

## Abstract

We describe LRSPAT (low-rank spatial toolkit), a fast and memory-efficient framework for approximating measures of spatial association for high-dimensional data. While LRSPAT can be applied to any multivariate spatial dataset, development was motivated by the computational challenge of identifying spatially variable genes in high-resolution spatial transcriptomics (ST) data generated by technologies such as 10x Visium HD, Xenium and Atera. LRSPAT leverages a truncated SVD of the expression data and a thresholded spatial weights matrix to perform reduced-rank reconstruction of spatial statistics in the quadratic form family, including global and local versions of Moran’s I, Geary’s C, and Getis-Ord G. A regularization approach is leveraged to account for the inflated null distribution of spatial statistics computed on latent variables. By performing key operations on the low-dimensional embeddings, LRSPAT is orders of magnitude faster than standard implementations with significantly lower memory requirements. Because the low-rank approach denoises and desparsifies ST data, LRSPAT is also more accurate than standard techniques at identifying genes with true spatial expression patterns. The dramatic improvements in execution time and memory consumption enable the genome-wide analysis of spatially variable genes (SVGs) and exploration of the full range of hyperparameters including spatial scale, distance metric, and embedding rank. This preprint outlines the background and mathematical details of the approach with limited preliminary results and a short conclusion.

## 1 Background

### 1.1 Spatial transcriptomics

Advances in spatial transcriptomics (ST) have revolutionized the study of complex tissues with high-resolution assays now able to capture the expression of thousands of genes at hundreds of thousands of tissue locations [1]. These techniques enable researchers to identify the cell types in the analyzed tissue, the phenotype of those cells, and, most importantly, how the cells are organized into interacting populations that drive tissue structure and function [2]. Two primary types of ST technologies are commercially available: high-throughput sequencing technologies (e.g., 10x Visium HD) and single-molecule fluorescent in situ hybridization or imaging-based technologies (e.g., 10x Xenium amd Atera). High-throughput sequencing ST assays capture mRNA transcripts in an untargeted fashion at multiple spots on a slide, which enables near transcriptome-wide profiling but has the downside that a direct mapping does not exist between spots and cells. Imaging ST assays, by contrast, use gene-specific probes (Xenium can target ∼5k genes and Atera ∼19k) to capture transcript abundance with subcellular resolution, which enables the assignment of reads to individual cells but has the downside that only the targeted genes are measured [3].

### 1.2 Spatially variable genes (SVGs)

A core analysis task for both types of ST data is the detection of spatially variable genes (SVGs). SVG detection methods fall under the more general class of univariate spatial association techniques, which quantify the degree of spatial autocorrelation and structural topology of a single variable distributed across an n-dimensional space. Many spatial association methods generate both local and global statistics, i.e., measures of spatial autocorrelation at each location and for the entire spatial dataset. Since the introduction of ST technologies, the field has seen a proliferation of methods designed to identify and quantify SVGs [4]. Existing approaches range from classic, non-parametric spatial autocorrelation metrics such as Moran’s I (MI) [5], Geary’s C (GC) [6], and Getis-Ord G (GOG) [7], to modern, ST-specific probabilistic frameworks like SPARK [8] and SPARK-X [9]. Recently, Yan et al. [4] categorized 34 state-of-the-art SVG algorithms according to three distinct biological signals: overall spatial variance, cell-type-specific variance, and spatial-domain markers. A recent benchmarking paper by Li et al. [10] evaluated a smaller group of 14 techniques. In this paper, we focus on a subset of univariate spatial statistics, the “quadratic form family”, that are mathematically based on a quadratic form involving the expression matrix **X** and either a spatial weights matrix **W** or graph Laplacian **L**, i.e., **X**^*T*^ **WX** or **X**^*T*^ **LX** (see Section 2.2 for full mathematical details).

### 1.3 Computational challenge of high-resolution spatial transcriptomics

Despite the abundance of SVG detection methods, systematic benchmarking by Li et al. [10] reveals severe operational limitations across the current state of the art [10]. Existing methodologies frequently suffer from highly inflated false-positive rates driven by extreme sparsity, dropout events, and zero-inflated technical noise inherent to single-cell and high-resolution ST platforms. Furthermore, traditional implementations of these spatial cross-correlation metrics require asymptotic time complexities of *O*(*N* ^2^) or *O*(*NP*) and necessitate loading very large dense matrices into RAM. These computational challenges have been exacerbated by the dramatic increase in spatial resolution for both sequencing-based (e.g., 10x Visium HD) and imaging-based (e.g., 10x Xenium, 10x Atera) ST platforms. The original Visium platform uses 5k 55 *µ*m diameter spots spaced 100 *µ*m apart that each contain 1-10 cells [11]. In contrast, Visium HD uses a continuous grid of ∼10.5 million 2×2 *µ*m squares [12] across each 6.5×6.5 mm capture area. Even at a coarsened resolution of 8 *µ*m, Visium HD can capture up to ∼625k tissue locations (∼950k for the planned Visium HD XL product). With sub-50 nm resolution, Xenium can match transcripts for ∼5k genes to individual cells across a 12×24 mm capture area and the just released Atera technology captures ∼19k human genes at single-cell resolution with almost 2x the capture area of Xenium.

For high-resolution ST platforms, exhaustive gene-level SVG analysis is computationally intractable for classic non-parametric methods such as MI, GC and GOG. To overcome this barrier, researchers typically limit SVG analysis to a subset of detected genes and use fast approximate methods such as SPARK-X. While this approach makes SVG detection computationally feasible, it ignores a large portion of the transcriptome and generates less accurrate SVG statistics. Importantly, Li et al. [10] demonstrated that classic quadratic form methods like MI frequently achieve greater accuracy than newer techiques with the added benefit of supporting both global and local versions, flexible distance metrics, and an approximate null distribution.

### 1.4 LRSPAT framework for accelerating the computation of quadratic form spatial statistics

To address these challenges, we developed the LRSPAT framework for the low-rank estimation of quadratic form spatial statistics including MI, GC and GOG. LRSPAT reconstructs gene-level spatial association statistics from statistics computed on a linear embedding of the ST data. By computing spatial association statistics on a low-dimensional embedding, LRSPAT offers an orders-of-magnitude improvement in execution time, dramatically lower memory requirements, and more accurrate SVG estimation via denoising property of reduced rank reconstruction. Importantly, the improved computational performance of LRSPAT enables genome-wide SVG analysis (vs. a focus on just a small subset of genes with high marginal variance) and the evaluation of spatial associations across a range of distance metrics, kernel shapes, and embedding ranks.

## 2 Methods

### 2.1 Notation

We assume a multivariate spatial dataset that captures *p* variables at *n* locations with measured values held in an *n* × *p* matrix **X**. The *n* measurements for a single variable, i.e, one column of **X**, are represented by the vector **x**. The standardized version of **x** (i.e., mean-centered and divided by the maximum likelihood estimate of the sample standard deviation) is represented by 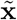 and the column-standardized version of version of **X** is represented by 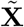. The Hadamard product (⊙) of **X** with itself, i.e., version where each element is squared, is represented by **X**^(2)^ = **X** ⊙ **X**.

An *n* × *m* matrix **S** holds the coordinates of each location in an m-dimensional space. Without loss of generality, we will assume these are x,y coordiantes in 2D Euclidean space. The coordinates in **S** are used to generate an *n* × *n* spatial weights matrix **W** with elements *w*_*i*,*j*_ that define the proximity or connection strength between locations *i* and *j*. Let *S*_0_ = ∑_*i*,*j*_ *w*_*ij*_. In our case, *w*_*i*,*j*_ will be inverse and exponentially scaled distances computed as 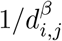 where *d*_*i*,*j*_ is an *L*_*p*_ distance, *β* is an exponential decay coefficient, and diagonal elements *d*_*i*,*i*_ = 0. In practice, **W** will also be sparsified using either a distance or k nearest neighbors (knn) threshold. Let 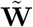 represent the row standardized version of **W**, i.e., 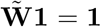 where **1** is a length *n* vector of ones.

### 2.2 Univariate spatial statistics and the quadratic form family

Univariate spatial statistics quantify the structural topology and degree of spatial autocorrelation for a given variable distributed across a spatial region. While traditional non-parametric univariate spatial statistics, such as Moran’s I (MI), Geary’s C (GC), and Getis-Ord G (GOG), are usually presented as summations over spatial neighbors, they can also be represented using a generalized quadratic form model. Specifically, these statistics quantify spatial autocorrelation by evaluating a target vector **x** ∈ ℝ^*n*^ (e.g., a column of **X**) against a predefined spatial structural operator **Ω** ∈ ℝ^*n*×*n*^. The unnormalized core of any statistic in this family reduces to the mathematical evaluation of a quadratic form:

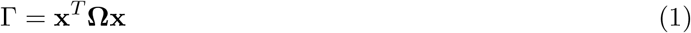

In a multivariate context, this is equivalent to the diagonal of **X**^*T*^ **ΩX**. By substituting different matrices for **Ω**, this model captures MI, GC and GOG, the three classic univariate spatial statistics supported by LRSPAT. For MI and GOG, spatial contiguity is quantified by setting **Ω** to the spatial weights matrix **W**. For GC, spatial dissimilarity is measured by setting **Ω** to the Graph Laplacian (**L** = **D** − **W**), where **D** is the diagonal degree matrix. While not explored in this paper, this quadratic form family includes classic bivariate spatial statistics such as Lee’s L [13], which replaces **Ω** by the squared spatial lag operator (**W**^*T*^ **W**). The quadratic form family also covers a number of more recent ST-specific methods including SPARK-X [9], Hotspot [14], and Giotto [15]. SPARK-X projects multiple spatial kernels against the ST gene expression matrix, which is equivalent to evaluating quadratic expressions equivalent to **X**^*T*^ **WX**, Hotspot computes local spatial autocorrelation using a localized Getis-Ord-like structure, and Giotto utilizes Moran’s I and Geary’s C statistics computed on a spatial nearest neighbor graph. While our low-rank framework could therefore be adapted to these specific techniques, we keep our focus on the classic MI, GC and GOG statistics in this paper.

Mathematical details for local and global versions of MI, GC and GOG are outlined in Sections 2.2.1, 2.2.2, and 2.2.3 below. Note that these mathematical definitions do not necessarily correspond to how the statistics would be calculated using numerically efficient code.

#### 2.2.1 Moran’s I

Global Moran’s I quantifies overall spatial clustering for a single variable using a mean-centered measure of spatial covariance. For a single standardized variable 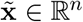 (i.e., a column of 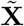), the statistic is defined by the standardized quadratic form:

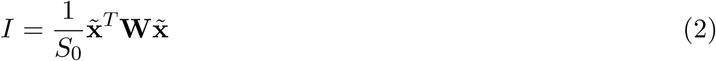

where *S*_0_ = ∑_*i*,*j*_ *w*_*ij*_. For the multivariate case, global MI statistics for the *p* variables held in a column-standardized matrix 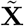 are computed as:

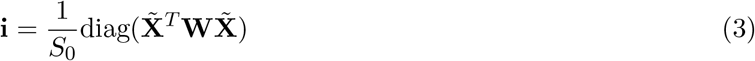

To identify localized spatial clusters or outliers (i.e., Local Indicators of Spatial Association, or LISA), Anselin’s Local Moran’s I values can be computed for all *p* variables at all *n* locations using the Hadamard product (⊙) between the standardized matrix 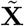 and its row-standardized spatial lag 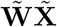:

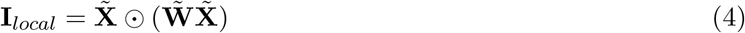

The average of all Local Moran’s I values for given variable (i.e, one column of **I**_*local*_) equals the global Moran’s I for that variable.

Statistical inference for Moran’s I evaluates the metric against a null hypothesis of spatial randomness. Because the expected value and variance under the analytical normality assumption depend exclusively on the sample size *n* and spatial weights matrix *W*, they act as scalar constants applied to generate analytical z-scores and p-values.

#### 2.2.2 Geary’s C

In contrast to Moran’s I, Geary’s C quantifies spatial dissimilarity by measuring squared differences between neighboring locations. This is defined mathematically using the unnormalized graph Laplacian matrix **L** = **D** − **W**, where **D** is the diagonal degree matrix (i.e., the row sums of **W**). For a single standardized variable **x**, Geary’s C is defined as:

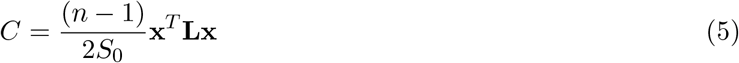

The Geary’s C values for all *p* variables are computed by evaluating the diagonal of the Laplacian cross-product:

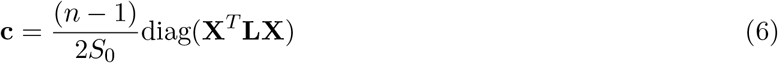

The local version of Geary’s C identifies individual locations with large squared differences from their immediate topological neighborhood. These are computed for all *p* variables at all *n* locations as:

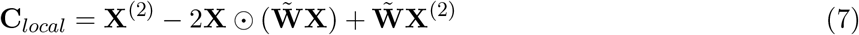

Note that this is defined using just the row-standardized weights matrix rather than the Laplacian for computational considerations.

Under the null hypothesis of spatial randomness, the expected value of Geary’s C is *E*[*C*] = 1.0. Under an assumption of normality, the variance depends strictly on the spatial weights and the number of locations, allowing for the computation of z-scores.

#### 2.2.3 Getis-Ord G

The Getis-Ord G (General G) statistic evaluates the spatial concentration of high or low values. Unlike Moran’s I, it does not rely on mean-centering; it operates strictly on non-negative raw values. For a single variable, it is defined as the spatial cross-product normalized by all possible non-spatial pairwise cross-products:

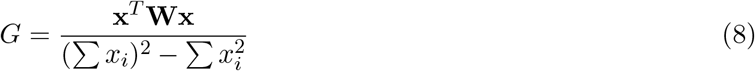

Extending to the full matrix **X**, the Getis-Ord G values for all *p* variables are defined as:

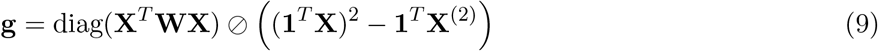

where ⊘ denotes element-wise division.

A local version of the Getis-Ord statistic 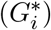 can be used to identify hotspots and coldspots. This is computed for all *p* variables and *n* locations as:

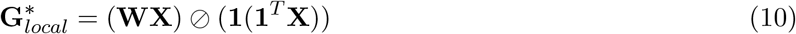

Because Getis-Ord G evaluates uncentered data, the exact analytical variance under a spatial randomization null hypothesis requires evaluation of the first through fourth moments of the raw data. This kurtosis mismatch frequently introduces significant numerical instability in high-sparsity ST regimes, a technical limitation that our low-rank approach specifically aims to regularize.

### 2.3 Low-rank approximation of quadratic form spatial statistics

Evaluating quadratic form univariate spatial statistics for an entire *n* × *p* expression matrix **X** requires extracting the diagonal elements of **X**^*T*^ **ΩX**. This has an asymptotic time complexity of *O*(*np*^2^) and necessitates loading massive, dense matrices into RAM. Since *n* typically exceeds 500k with *p* between 5-20k for high-resolution spatial transcriptomics (e.g., Visium HD, Xenium 5k, and Atera), this type of direct spatial analysis is computationally intractable for modern ST datasets without dramatic reductions in *p* (via feature selection) and/or *n* (via sketching).

To address this challenge without discarding genes or locations, we explored low-rank approximations of global and local versions of the MI, GC and GOG statistics detailed above. Specifically, by applying a truncated singular value decomposition (SVD) (as computed by an efficient randomized [16] or iterative [17] algorithm) to either the raw or mean-centered expression data (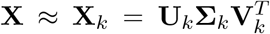 or 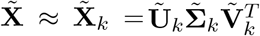 with rank *k* ≪ *n, p*), we can isolate the computation to the dense latent embeddings (**U**_*k*_ ∈ ℝ^*n*×*k*^) and only evaluate a compact *k* × *k* latent spatial covariance matrix:

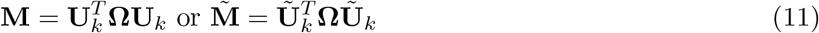

We will use a subscript to represent the specific value of **Ω**, e.g., **M**_**W**_ when **Ω** = **W**.

This low-rank approach bounds memory usage to the latent space and reduces the asymptotic time complexity to *O*(*ek*), where *e* is the number of non-zero edges in the sparse spatial network **Ω**, e.g., number of non-zero weights in **W**. This low-rank framework not only provides a dramatically lower computational cost but also acts as a powerful structural regularizer of the biological signal. Specifically, restricting the evaluation of **X**^*T*^ **ΩX** to a rank *k* embedding filters out stochastic noise and mitigates the impact of sparsity. This structural shrinkage prevents the false-positive inflation observed in standard implementations and rescues low magnitude spatial test statistics from technical dropout.

It is important to note that this low-rank approach yields identical results to those generated by analyzing the quadratic form for the rank *k* reconstruction 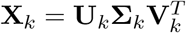:

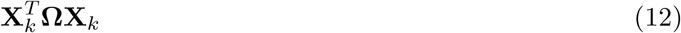

While naively reconstructing the full **X**_*k*_ would still gain the denoising/regularization benefits, it would be unhelpful from a computational complexity standpoint. If the target **X** is already dense, it would incur the additional cost of computing the truncated SVD and evaluating 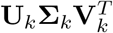. For highly sparse data, e.g., spatial transcriptomics, the impact would be catastrophic since it replaces analysis of a very sparse **X** with analysis of a dense **X**_*k*_. The computational benefits of an SVD-based low-rank approach are obtained by restricting the expensive analyses to just the rank *k* latent space and the *k* × *k* latest spatial convariance matrix **M**.

On the computational front, it is also important to note that, for high-resolution ST data, mean centering **X** will turn a large but very sparse matrix into a dense matrix with a dramatic impact on memory requirements and execution time. Standard ST analysis pipelines avoid this problem by only performing truncated PCA (i.e., SVD on the mean-centered matrix) after feature selection. In the remainder of this paper, we ignore this issue and assume that a truncated SVD can be computed effectively on either **X** or, for Moran’s I, on 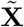. In practice, mean-centering can either be skipped on ST data given the very low mean values of most genes or performed only for a small fraction of genes with a large mean.

Sections 2.3.1, 2.3.2, and 2.3.3 below detail how this low-rank framework can be leveraged to approximate global and local versions of MI, GC, and GOG. As noted above, these mathematical definitions do not necessarily correspond to how the computations would actually be implemented using efficient code, i.e., the computations can be limited to the rank *k* latent space without requiring reconstruction of a full *n* × *p* matrix.

#### 2.3.1 Low-rank Moran’s I

We can compute a rank *k* approximation of the global Moran’s I statistics (**i**_*lr*_) for all *p* variables using the *k* × *k* latent spatial covariance matrix 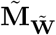 as defined in (11) and the variable loadings from the truncated 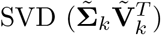:

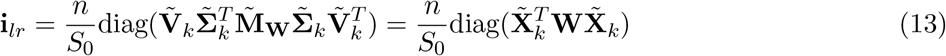

As detailed above, computing 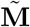 requires only *O*(*ek*) time, and the subsequent projection 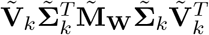 reduces the feature-space evaluation from *O*(*np*^2^) to *O*(*k*^2^*p*). Because *k* ≪ *n, p*, this yields a massive reduction in execution time while also bounding memory allocation to the *k* × *k* and *k* × *p* latent spaces.

To compute low-rank approximations of local Moran’s I statistics, we first reconstruct the matrix of spatially lagged embeddings, 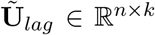, by directly multiplying the row-standardized weights matrix 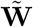 against the embeddings:

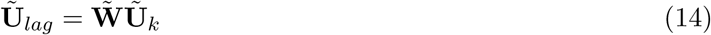

Given **Ũ**_*lag*_, low-rank approximations of the local Moran’s I values for all *p* variables at all *n* locations is generated via the Hadamard product (⊙) of the reconstructed data and its corresponding reconstructed spatial lag:

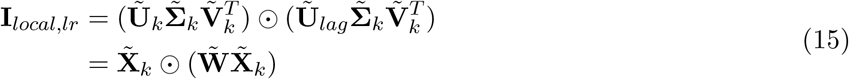

#### 2.3.2 Low-rank Geary’s C

We can compute rank *k* estimates of the global Geary’s C values (**c**_*lr*_) for all *p* variables using the *k* × *k* latent spatial covariance matrix **M**_**L**_ as defined in (11) and the variable loadings from the truncated SVD 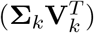:

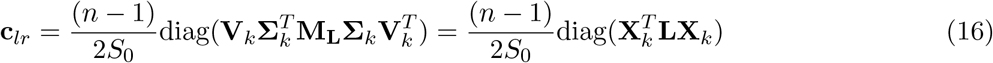

Low-rank estimates of the local Geary’s C values can be computed for the entire dataset as:

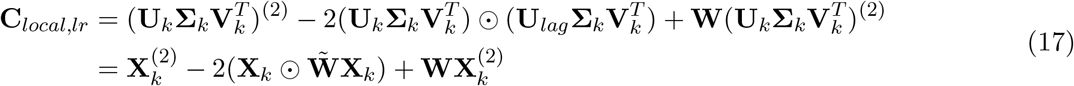

#### 2.3.3 Low-rank Getis-Ord G

The vector of rank *k* reconstructed of global Getis-Ord G statistics (**g**_*lr*_) is computed using the *k* × *k* latent spatial covariance matrix **M**_**W**_:

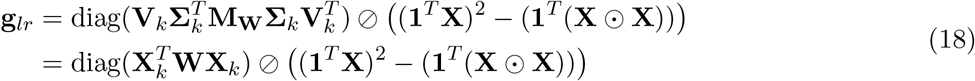

Note that this formulation uses the raw **X** to form the denominator rather than a low-rank reconstruction. This is done to prevent the shrinkage of this normalizing term that would result from an approximation based on a truncated SVD and the significant distortion this would cause to estimates of the analytical variance of the GOG statistics under the null of spatial randomization.

Low rank approximations of the local Getis-Ord *G*^∗^ statistics can be used to identify biological hot/cold spots without being confounded by technical noise/sparsity. These are computed as:

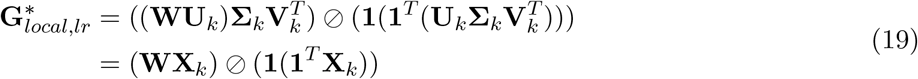

### 2.4 Bayesian regularization of latent spatial variance

While projecting high-dimensional spatial data into a low-rank latent space provides both denoising and a significant computational acceleration, it introduces a problematic mathematical artifact. By definition, truncated matrix decompositions (e.g., SVD) capture the dominant axes of variance within a dataset. Because true variance for experimental data (e.g., spatial transcriptomics) is spatially organized (e.g., distinct cell types, morphological domains), the resulting latent embeddings (**U**_*K*_) will usually have significant spatial autocorrelation. This is problematic when evaluating non-spatial noise variables reconstructed from these embeddings. Even if a variable represents pure stochastic noise, projecting it into the latent space forces it to adopt a linear combination of the embeddings that each have a large spatial association. Because standard non-parametric spatial statistics (like Moran’s I or Geary’s C) are mathematically scale-invariant, they normalize the spatial cross-product by the marginal variance. Consequently, the tiny latent magnitude of a noise variable allows it to artificially inherit the massive spatial associations of the embeddings. Left uncorrected, this structural artifact will generate a high false-positive rate for noise variables with low empirical variance.

To correct this artifact, we must break the scale-invariance of the standard spatial estimators. We propose an Empirical Bayesian [18] shrinkage framework that explicitly penalizes variables that lack sufficient absolute variance in the rank-*k* space, preventing them from adopting the spatial structure of the dominant embeddings. Specifically, we estimate a penalty, *ϵ*, derived from the distribution of latent variances. We apply this Bayesian prior directly to the variance denominator of our generalized multivariate quadratic form (*γ*):

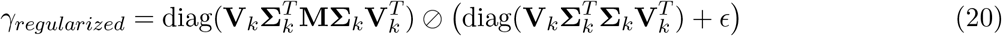

We estimate *ϵ* as a quantile-based threshold of the distribution of variances of the *k* latent variables. By setting the penalty to a high quantile (e.g., the 90th percentile of latent variance, 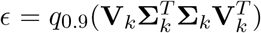, we can retain true spatial variables and suppress statistics for non-spatial variables below the significance threshold. Variables with strong spatial structure possess large latent variance that easily overpowers the *ϵ* penalty and thus retain significant *z*-scores. By contrast, purely stochastic variables project poorly into the latent space, resulting in tiny latent space variance. For these variables, the *ϵ* penalty dominates the denominator expression, shrinking the inherited spatial cross-product toward the null expectation of zero.

## 3 Results

### 3.1 Execution time and memory use

The following table details the theoretical asymptotic computational time and memory requirements for standard and low-rank versions of global and local Moran’s I, Geary’s C and Getis-Ord G. As detailed in Section 2.1, *n* represents the number of spatial location, *p* the number of variables measured at each location, *k* the low-dimensional rank with *k* ≪ *n, p*, and *e* the number of non-zero elements in the sparse spatial weights matrix **W**.

**Table 1:** Asymptotic time and memory complexities for standard vs. low-rank spatial statistics.

| Statistic | Type | Execution time |  | Peak RAM |  |
| --- | --- | --- | --- | --- | --- |
|  |  | Standard | Low-rank | Standard | Low-rank |
| **Moran’s I** | Global | $O(pe)$ | $O(ke + pk^2)$ | $O(e + np)$ | $O(e + nk + pk)$ |
| | Local (LISA) | $O(pe)$ | $O(ke + npk)$ | $O(e + np)$ | $O(e + nk + pk)^\dagger$ |
| **Geary’s C** | Global | $O(pe)$ | $O(ke + pk^2)$ | $O(e + np)$ | $O(e + nk + pk)$ |
| | Local | $O(pe)$ | $O(ke + npk)$ | $O(e + np)$ | $O(e + nk + pk)^\dagger$ |
| **Getis-Ord G** | Global | $O(pe)$ | $O(ke + pk^2 + np)$ | $O(e + np)$ | $O(e + np + pk)$ |
| | Local ($G_i^*$) | $O(pe)$ | $O(ke + npk)$ | $O(e + np)$ | $O(e + nk + pk)^\dagger$ |

<sup>†</sup> Denotes memory footprint when utilizing a factorized representation (storing the decoupled embedding and coordinate matrices). Explicitly instantiating the full dense local statistic output requires $O(np)$ RAM.

Standard global evaluations force repeated spatial network traversals for every variable, scaling at *O*(*pe*). By projecting the topology into the rank-*k* manifold, the graph traversal is computed just once (*O*(*ke*)), reducing the *p*-dependent scaling to simple vector operations bounded by *O*(*pk*^2^). Traditional local statistics allocate multiple heavy dense matrices during intermediate cross-product lags, bottlenecking RAM at *O*(*np*). By enforcing localized computations inside the reduced *k*-dimensional space, we execute dense-dense operations that bypass allocating full raw spatial lag arrays entirely, restricting overhead to the *nk* and *kp* latent spaces. Note that the low-rank global Getis-Ord G incorporates an additional *O*(*np*) factor in both time and memory. This reflects our targeted hybrid formulation that evaluates the normalization denominator strictly over the raw, uncentered **X** matrix to compute the analytical expected value and prevent denominator drift from least-squares truncation.

### 3.2 Detection of spatially-variable genes

In addition to the significant reductions in execution time and memory cost detailed above, the LRSPAT framework can more effectively identify variables with a true spatial association when the target data is high-dimensional, sparse and noisy, characteristics that are exemplified by high-resolution ST data. To evalute the denoising performance of LRSPAT, we explored both simulated ST data with a known ground truth and Visium HD data for a mouse brain coronal section. For both the simulated and real data:

- A rank 30 truncated SVD was computed on the data matrix **X** using the randomized algorithm implemented by the *rsvd* R package [16].
- The spatial weights matrix **W** was computed using inverse-squared Euclidean distances that were thresholded at a Euclidean distance that retained an average of 20 neighbors. To ensure computational efficiency, this was implemented using the KD-tree logic in the *dbscan* R package [19].
- Standard and low-rank versions of global MI, GC and GOG statistics (as defined in Section 2 with the low-rank regularization *ϵ* set to the 0.2 quantile) were computed on **W** and either **X** or the output of the truncated SVD.

For the simulation study, the data was generated to match the characteristics of ST data with n=5000, p=5000, k=30, 95% sparsity, and 10% of the genes with a true spatial association. Figure 1 visualizes the standard and low-rank spatial statistics a single simulated dataset. In this case, the low-rank approach generates lower AUC-ROC and PR-AUC values for all three global statistics validating the expected denoising performance.

**Figure 1.**
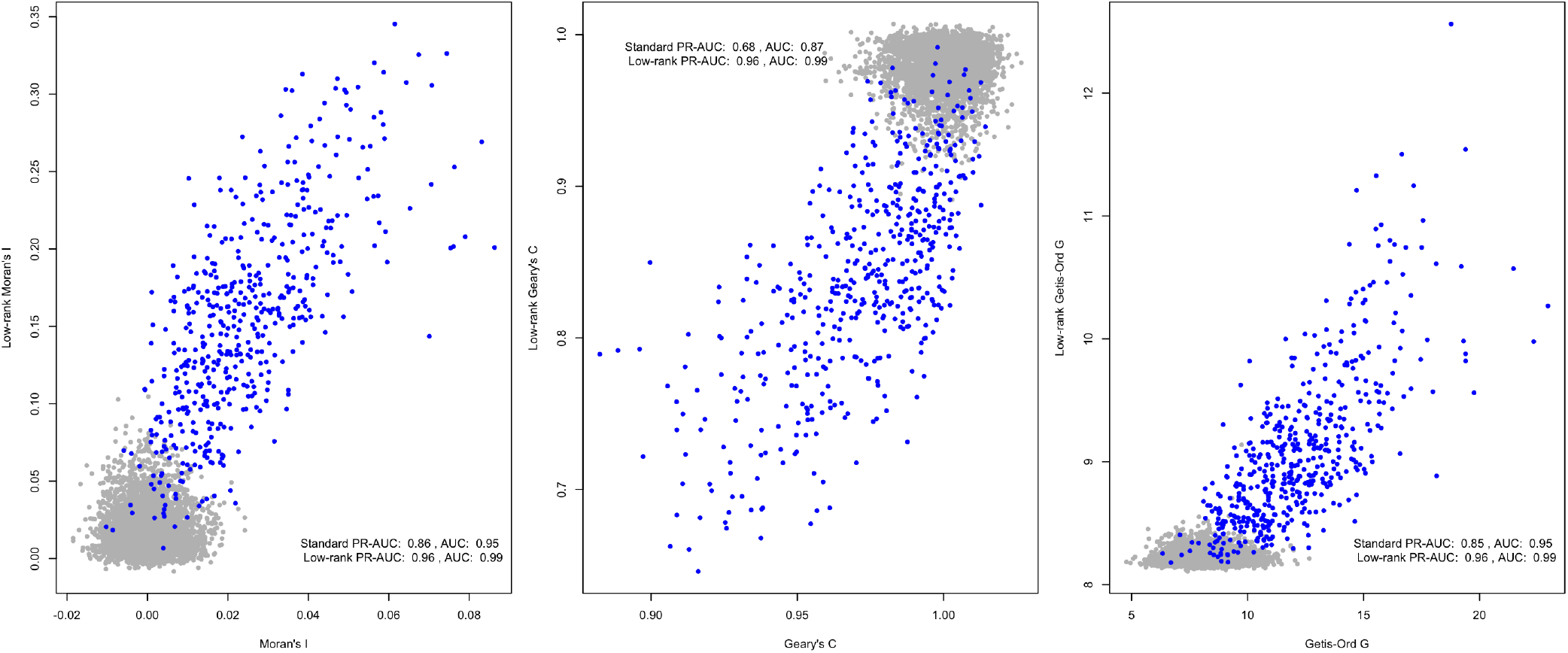
Detection of variables with a true spatial association in simulated data according to standard and low-rank global Moran’s I, Geary’s C, and Getis-Ord G. Variables with a true association are shown in blue and those without in grey.

For evaluation on real ST data, we used a mouse brain Visium HD dataset for an FFPE coronal section (available from 10x at https://www.10xgenomics.com/datasets/visium-hd-cytassist-gene-expression-mouse-brain-fresh-frozen. The data was processed using the v5.3.0 of the Seurat [20] framework at 16 *µ*m resolution with log-normalization and just the top 2k highly-variable genes (HVGs) retained in **X** to match standard processing logic. Figure 2 visualizes the standard and low-rank spatial statistics for this Visium data. The Allen Mouse Brain Atlas list of 500 genes with brain region-specific expression (available from “https://alleninstitute.github.io/abc_atlas_access/_downloads/54f8c94c59bb7c322c5f933e0b151681/gene_list.html, as curated in Yao et al. [21]) was used as the group of genes with an assumed true spatial association for computing AUC values (note that only 247 of these genes were among the top 2k HVGs). Not surprisingly, all AUC values computed on the real data are much lower than for the simulate data, which is consistent with the approximate nature of the ground truth and the strong spatial signature used in the simulation model. Similar to the simulation results, the low-rank version of MI and GC again produce higher ROC-AUC and PR-AUC than the standard implementations. For GOG, the AUC values are slightly higher for the standard implementation.

**Figure 2.**
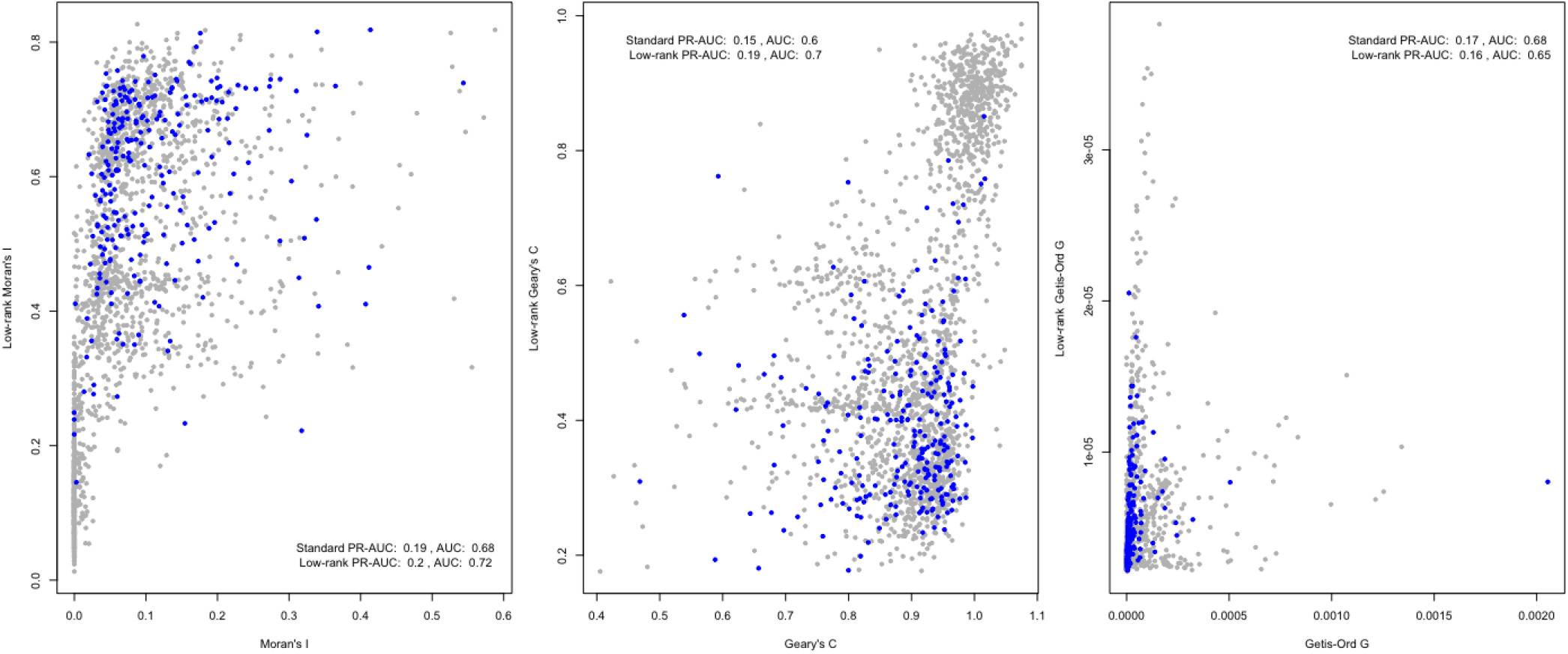
Detection of genes from the Allen Mouse Brain Atlas list of 500 genes with brain region-specific expression according to standard and low-rank global Moran’s I, Geary’s C, and Getis-Ord G. Genes from the Allen Brain Atlas are shown in blue with other genes in grey.

## 4 Conclusion

Exhaustive spatial association analysis for modern, high-resolution spatial transcriptomics is frequently derailed by the dual bottlenecks of high computational cost and severe zero-inflated technical noise. In this preprint, we introduced LRSPAT, a computational framework that projects univariate spatial statistics of the quadratic form family into an orthogonal latent space. By isolating the spatial statistic calculation to a compact *k* × *k* latent spatial covariance matrix, LRSPAT drastically reduces standard dense execution times to *O*(*k*^2^*p*) and strictly bounds peak memory consumption, rendering whole-transcriptome spatial mapping computationally tractable for massive coordinate systems. Beyond mere computational acceleration, our application of an Empirical Bayesian shrinkage prior acts as a critical structural regularizer. By neutralizing the scale-invariant artifact inherent to low-rank reconstructions, LRSPAT successfully performs manifold denoising, recovering the true structural topology of spatially variable genes that are otherwise obscured by technical dropout in standard non-parametric implementations.

## References

[1] Williams, C.G., Lee, H.J., Asatsuma, T., Vento-Tormo, R., Haque, A.: An introduction to spatial transcriptomics for biomedical research. Genome Med 14(1), 68 (2022). doi:10.1186/s13073-022-01075-1

[2] Tian, L., Chen, F., Macosko, E.Z.: The expanding vistas of spatial transcriptomics. Nat Biotechnol 41(6), 773–782 (2023). doi:10.1038/s41587-022-01448-2

[3] Marco Salas, S., Kuemmerle, L.B., Mattsson-Langseth, C., Tismeyer, S., Avenel, C., Hu, T., Rehman, H., Grillo, M., Czarnewski, P., Helgadottir, S., Tiklova, K., Andersson, A., Rafati, N., Chatzinikolaou, M., Theis, F.J., Luecken, M.D., Wählby, C., Ishaque, N., Nilsson, M.: Optimizing xenium in situ data utility by quality assessment and best-practice analysis workflows. Nat Methods 22(4), 813–823 (2025). doi:10.1038/s41592-025-02617-2

[4] Yan, G., Hua, S.H., Li, J.J.: Categorization of 34 computational methods to detect spatially variable genes from spatially resolved transcriptomics data. Nat Commun 16(1), 1141 (2025). doi:10.1038/s41467-025-56080-w

[5] Moran, P.A.P.: A test for the serial independence of residuals. Biometrika 37(1-2), 178–81 (1950)

[6] Geary, R.C.: The contiguity ratio and statistical mapping. The Incorporated Statistician 5, 115 (1954). doi:10.2307/2986645

[7] Getis, A., Ord, J.K.: The analysis of spatial association by use of distance statistics. Geographical Analysis 24(3), 189–206 (1992). doi:10.1111/j.1538-4632.1992.tb00261.x

[8] Sun, S., Zhu, J., Zhou, X.: Statistical analysis of spatial expression patterns for spatially resolved transcriptomic studies. Nat Methods 17(2), 193–200 (2020). doi:10.1038/s41592-019-0701-7

[9] Zhu, J., Sun, S., Zhou, X.: Spark-x: non-parametric modeling enables scalable and robust detection of spatial expression patterns for large spatial transcriptomic studies. Genome Biol 22(1), 184 (2021). doi:10.1186/s13059-021-02404-0

[10] Li, Z. M Patel, Z., Song, D., Yasa, S.N., Cannoodt, R., Yan, G., Li, J.J., Pinello, L.: Systematic benchmarking of computational methods to identify spatially variable genes. Genome Biol 26(1), 285 (2025). doi:10.1186/s13059-025-03731-2

[11] Ståhl, P.L., Salmén, F., Vickovic, S., Lundmark, A., Navarro, J.F., Magnusson, J., Giacomello, S., Asp, M., Westholm, J.O., Huss, M., Mollbrink, A., Linnarsson, S., Codeluppi, S., Borg, Å., Pontén, F., Costea, P.I., Sahlén, P., Mulder, J., Bergmann, O., Lundeberg, J., Frisén, J.: Visualization and analysis of gene expression in tissue sections by spatial transcriptomics. Science 353(6294), 78–82 (2016). doi:10.1126/science.aaf2403

[12] Polański, K., Bartolomé-Casado, R., Sarropoulos, I., Xu, C., England, N., Jahnsen, F.L., Teichmann, S.A., Yayon, N.: Bin2cell reconstructs cells from high resolution visium hd data. Bioinformatics 40(9), 546 (2024). doi:10.1093/bioinformatics/btae546

[13] Lee, S.-I.: Developing a bivariate spatial association measure: An integration of pearson’s r and moran’s i. Journal of Geographical Systems 3(4), 369–385 (2001)

[14] DeTomaso, D., Yosef, N.: Hotspot identifies informative gene modules across modalities of single-cell genomics. Cell Systems 12(5), 438–450 (2021). doi:10.1016/j.cels.2021.04.004

[15] Dries, R., Zhu, Q., Dong, R., Eng, C.-H.L., Li, H., Liu, K., Fu, Y., Zhao, T., Sarkar, A., Bao, F., George, R.E., Pierson, N., Cai, L., Yuan, G.-C.: Giotto: a toolbox for integrative analysis and visualization of spatial expression data. Genome Biology 22(1), 78 (2021). doi:10.1186/s13059-021-02286-2

[16] Erichson, N.B., Voronin, S., Brunton, S.L., Kutz, J.N.: Randomized matrix decompositions using r. \ Journal of Statistical Software 89(11), 1–48 (2019). doi:10.18637/jss.v089.i11

[17] Baglama, J., Reichel, L., Lewis, B.W.: Irlba: Fast Truncated Singular Value Decomposition and Principal Components Analysis for Large Dense and Sparse Matrices. (2022). R package version 2.3.5.1. https://CRAN.R-project.org/package=irlba

[18] Ritchie, M.E., Phipson, B., Wu, D., Hu, Y., Law, C.W., Shi, W., Smyth, G.K.: limma powers differential expression analyses for rna-sequencing and microarray studies. Nucleic Acids Res 43(7), 47 (2015). doi:10.1093/nar/gkv007

[19] Hahsler, M., Piekenbrock, M., Doran, D.: dbscan: Fast density-based clustering with R. Journal of Statistical Software 91(1), 1–30 (2019). doi:10.18637/jss.v091.i01

[20] Butler, A., Hoffman, P., Smibert, P., Papalexi, E., Satija, R.: Integrating single-cell transcriptomic data across different conditions, technologies, and species. Nat Biotechnol 36(5), 411–420 (2018). doi:10.1038/nbt.4096

[21] Yao, Z., van Velthoven, C.T.J., Kunst, M., Zhang, M., McMillen, D., Lee, C., Jung, W., Goldy, J., Abdelhak, A., Aitken, M., Baker, K., Baker, P., Barkan, E., Bertagnolli, D., Bhandiwad, A., Bielstein, C., Bishwakarma, P., Campos, J., Carey, D., Casper, T., Chakka, A.B., Chakrabarty, R., Chavan, S., Chen, M., Clark, M., Close, J., Crichton, K., Daniel, S., DiValentin, P., Dolbeare, T., Ellingwood, L., Fiabane, E., Fliss, T., Gee, J., Gerstenberger, J., Glandon, A., Gloe, J., Gould, J., Gray, J., Guilford, N., Guzman, J., Hirschstein, D., Ho, W., Hooper, M., Huang, M., Hupp, M., Jin, K., Kroll, M., Lathia, K., Leon, A., Li, S., Long, B., Madigan, Z., Malloy, J., Malone, J., Maltzer, Z., Martin, N., McCue, R., McGinty, R., Mei, N., Melchor, J., Meyerdierks, E., Mollenkopf, T., Moonsman, S., Nguyen, T.N., Otto, S., Pham, T., Rimorin, C., Ruiz, A., Sanchez, R., Sawyer, L., Shapovalova, N., Shepard, N., Slaughterbeck, C., Sulc, J., Tieu, M., Torkelson, A., Tung, H., Valera Cuevas, N., Vance, S., Wadhwani, K., Ward, K., Levi, B., Farrell, C., Young, R., Staats, B., Wang, M.-Q.M., Thompson, C.L., Mufti, S., Pagan, C.M., Kruse, L., Dee, N., Sunkin, S.M., Esposito, L., Hawrylycz, M.J., Waters, J., Ng, L., Smith, K., Tasic, B., Zhuang, X., Zeng, H.: A high-resolution transcriptomic and spatial atlas of cell types in the whole mouse brain. Nature 624(7991), 317–332 (2023). doi:10.1038/s41586-023-06812-z

